# Polygenic selection drives the evolution of convergent transcriptomic landscapes across continents within a Nearctic sister-species complex

**DOI:** 10.1101/311464

**Authors:** Clément Rougeux, Pierre-Alexandre Gagnaire, Kim Praebel, Ole Seehausen, Louis Bernatchez

## Abstract

In contrast to the plethora of studies focusing on the genomic basis of adaptive phenotypic divergence, the role of gene expression during speciation has been much less investigated and consequently, less understood. Yet, the convergence of differential gene expression patterns between closely related species-pairs might reflect the role of natural selection during the process of ecological speciation. Here, we test for intercontinental convergence in differential transcriptional signatures between limnetic and benthic sympatric species-pairs of Lake Whitefish (*Coregonus clupeaformis*) and its sister-lineage, the European Whitefish (*C. lavaretus*), using six replicated sympatric species-pairs (two in North America, two in Norway and two in Switzerland). We characterized both sequence variation in transcribed regions and differential gene expression between sympatric limnetic and benthic species across regions and continents. Our first finding was that differentially expressed genes (DEG) between limnetic and benthic whitefish tend to be enriched in shared polymorphism among sister-lineages. We then used both genotypes and co-variation in expression in order to infer polygenic selection at the gene level. We identified parallel outliers and DEG involving genes primarily over-expressed in limnetic species relative to the benthic species. Our analysis finally revealed the existence of shared genomic bases underlying parallel differential expression across replicated species pairs from both continents, such as a *cis*-eQTL affecting the pyruvate kinase expression level involved in glycolysis. Our results are consistent with a longstanding role of natural selection in maintaining transcontinental diversity at phenotypic traits involved in ecological speciation between limnetic and benthic whitefishes.

## INTRODUCTION

Deciphering the genomic basis of differential adaptations between divergent populations, ultimately leading to ecological speciation, has been of foremost interest over the last decade. Adaptive divergence implies different population phenotypic responses to constraints associated with selective pressures stemming from different environments. This is particularly well illustrated by the occurrence of independent parallel phenotypic evolution among closely related and locally adapted nascent species (Endler 1986; Orr 2005; Losos 2011). Parallel phenotypic evolution can emerge from repeated divergence of the same genomic regions (Conte et al. 2012) or from different genes involved in similar or different biological pathways (Cohan & Hoffmann 1989; Losos 2011), and has been associated with changes in gene expression during adaptive divergence (Pavey et al. 2010; Manceau et al. 2011; Harrison et al. 2012). Genetic variation underlying parallel phenotypic changes may originate from parallelism at the molecular level that has arisen from *de novo* mutations affecting the same genes (Manceau et al. 2010; Rockman 2012). However, such mutations are generally associated with loci of large effect controlling the expression of a given phenotypic trait with a mono-/oligo-genic architecture (Manceau et al. 2010). This contrasts with the polygenic architecture of most complex traits, including those thought to be involved in ecological speciation, and more generally in adaptation (Savolainen et al. 2013; Yeaman 2015; Gagnaire & Gaggiotti 2016). For such traits, standing genetic variation is usually seen as an important source of adaptive mutations (Welch & Jiggins 2014). Several recent studies have showed that standing variation may originate from past admixture events (Roesti et al. 2014; Martin et al. 2015; Meier et al. 2017; Rougeux et al. 2017), suggesting an important role of anciently diverged variants in the process of ecological speciation (Marques *et al.* 2019). Despite the increasing number of studies underlining the fundamental role of standing variation as the main fuel for adaptation (Barrett & Schluter 2008; Schrider & Kern 2017), relatively few have focused on the possible consequences of different levels of standing genetic variation (*e.g.*, because of different historical contingencies) across populations on the fate of parallel phenotypic evolution (Nelson & Cresko 2018).

In this study, we compare the genomic basis of limnetic-benthic divergence among sympatric species-pairs from two different evolutionary lineages: the North American lake whitefish (*Coregonus clupeaformis* species complex) and the European whitefish (*C. lavaretus* species complex). The two sister-lineages *C. clupeaformis* (from North America) and *C. lavaretus* (from Europe) became geographically isolated ∼500,000 years ago and have evolved independently since then (Bernatchez & Dodson 1991; 1994; Jacobsen et al. 2012). In both lineages, several isolated lakes harbour partially reproductively isolated sympatric benthic (normal) and limnetic (dwarf) species-pairs. Limnetic and benthic sympatric species-pairs display sufficient levels of reproductive isolation to be considered as valid biological species (Kottelat & Freyhof 2007). Therefore, we will use the term “species” throughout the manuscript to refer strictly to limnetic and benthic whitefish, independently of their continent of origin. This system thus offers a valuable model to study the genomic and transcriptomic underpinnings of parallel differential adaptations leading to ecological speciation in independent lineages. The European whitefish species-pairs appear to be the result of a secondary contact between glacial sub-lineages (Rougeux et al. 2019), resulting in intra-lacustrine evolution of benthic and limnetic species across Scandinavian and Alpine lakes (Douglas et al. 1999; Østbye et al. 2005; 2006). The North American lake whitefish sympatric species-pairs are also the result of a post-glacial secondary-contact between two glacial sub-lineages during the late Pleistocene. In both lineages, the allopatric phase that likely lasted about 60,000-100,000 years has allowed the accumulation of genomic incompatibilities between sub-lineages, while secondary contact around 12,000 years ago has provoked character displacement in sympatry, leading to the current phenotypic and ecological divergence (Bernatchez & Dodson 1990; 1991; Pigeon et al. 1997; Rougeux et al. 2017).

Lake whitefish have been the subject of numerous studies pertaining to the ecological and genomic basis of adaptive divergence between limnetic and benthic species. The limnetic species differ from the benthic species in their use of habitat and trophic resources, with a higher metabolic rate and more active swimming behaviour for foraging and predator avoidance (Bernatchez et al. 1999; Trudel et al. 2001; Rogers et al. 2002), reduced energy allocated to growth relative to benthic whitefish (Trudel et al. 2001; Rogers & Bernatchez 2004) and differences in morphology, life history and physiological traits (Rogers & Bernatchez 2007; Dalziel et al. 2015; 2017b), which are under polygenic control (Gagnaire et al. 2013b; Laporte et al. 2015). Given the recent divergence between American and European sister lineages, an important part of the divergence between limnetic and benthic would have predated the divergence between continents, especially for genes associated to phenotypic diversification. Thus, the divergence between species may have stemmed from divergent selection acting on standing genetic variation. A possible outcome could be that divergent selection between species has been actively maintaining shared polymorphism at selected variants across continents, protecting them from being lost by drift.

The main goal of this study was to investigate the role of ancestral polymorphism on differential transcriptional signatures between limnetic and benthic species. Our aim was to test the general hypothesis that an overlapping polygenic basis underlies the parallel phenotypic divergence observed between sympatric species-pairs from sister lineages living on two different continents. Specifically, i) we first documented the amount and functional role of shared ancestral genetic polymorphism within coding genes among populations of the entire system, ii) we then quantified the extent of differential gene expression between benthic and limnetic species at the local (lake), regional and inter-continental scales, iii) we tested whether genes differentially expressed display an excess of shared polymorphism between sister-lineages, and finally iv) explored associations between polymorphism and variation in expression at the gene level.

## MATERIALS AND METHODS

### Sample collection, library preparation and sequencing

*C. clupeaformis* samples were collected from Indian Lake and Cliff Lake, Maine (USA) (Fig 1) in 2010. These lakes are part of a well-studied lake whitefish system (Bernatchez et al. 2010) and comprise the most divergent species-pairs along the divergence continuum described in previous studies (Renaut et al. 2012; Gagnaire et al. 2013b; Rougeux et al. 2017). In parallel, *C. lavaretus* individuals were sampled in two Scandinavian lakes in Norway (2014): Skrukkebukta, Langfjordvatn and two alpine lakes in Switzerland (2012): Zurich and Lucerne (Fig 1). We chose these European lakes as they each contained only two sympatric limnetic-benthic populations (*i.e.*, excluding potential genetic interactions with other sympatric whitefish forms that occur in other lakes) consistent with our sampling for *C. clupeaformis*. For each species pair, six benthic and six limnetic individuals were sampled (72 samples in total). Fresh liver biopsies were taken, flash frozen, and stored at −80°C for Lake whitefish, while European whitefish livers were stored directly in RNAlater. All individuals were sampled during summer for each locality, and only mature males were selected for this study in order to reduce sex-specific and life-stage gene expression variability.

**Fig 1.**
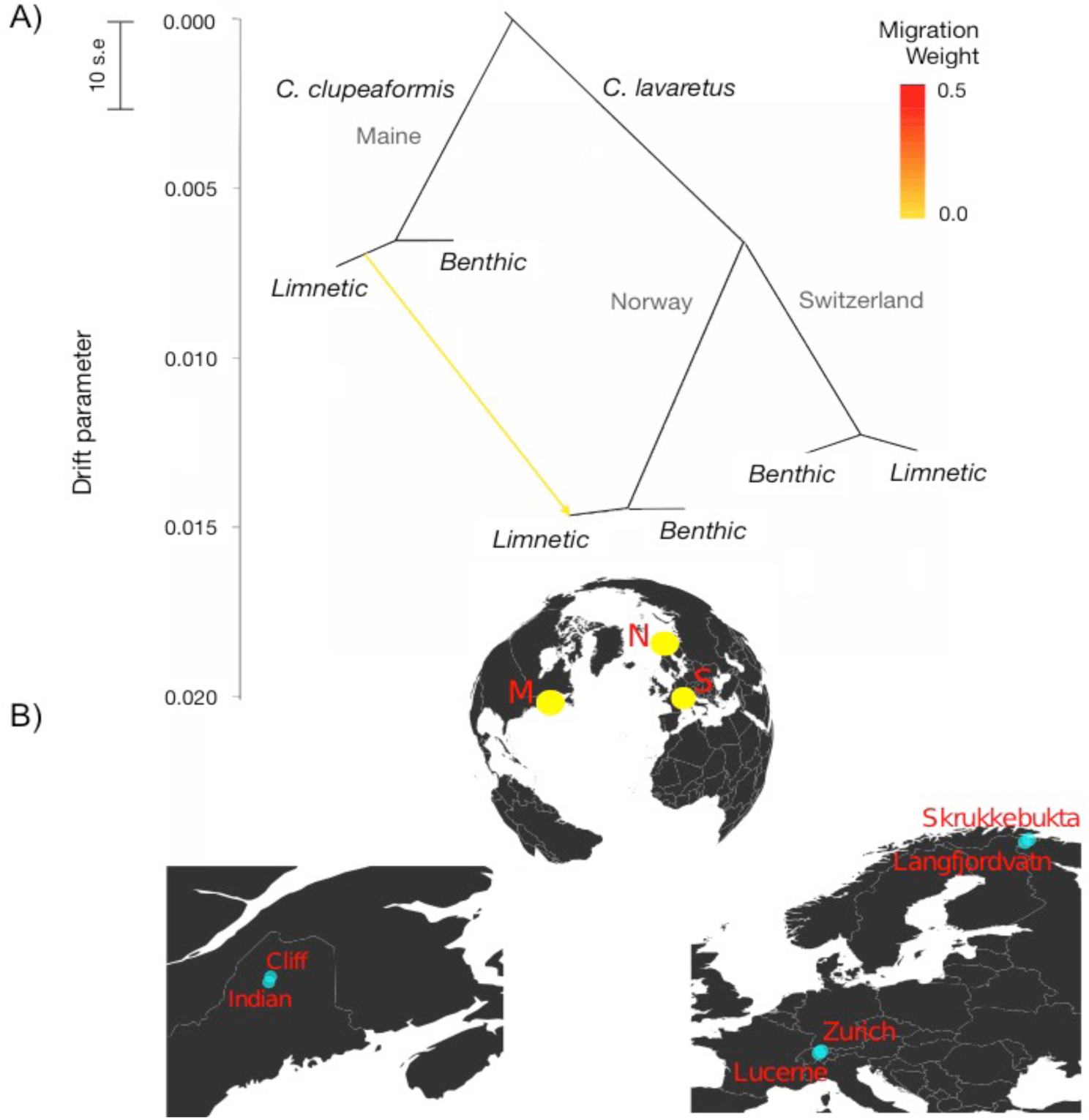
Details about the whitefish study system. A) Treemix analysis illustrating independent differentiation between sympatric Benthic and Limnetic species, from the closely related sister lineages *C. clupeaformis* in North America and *C. lavaretus* in Europe. The lake level was removed by merging populations from the same region for more clarity. Vertical branch lengths are proportional to the amount of genetic drift in each branch, the scale bar indicates 10 times the average standard error (s.e) of the entries in the covariance matrix between pairs of populations. The color scale indicates the weight of inferred migration events or shared ancestral polymorphism in absence of possible gene flow across continents, represented by the arrow between tree tips. B) Locations of the three sampled regions (yellow circles) M: Maine for *C. clupeaformis*, N: Norway and S: Switzerland for *C. lavaretus*. Two lakes (blue circles) containing sympatric species-pairs were sampled per region.

Prior to extraction, all samples were assigned randomly to different extraction groups (*i.e.*, groups of samples extracted at the same time) to minimise batch effect of any specific group of samples. Total RNA was extracted from liver tissue pieces of equal size using the RNAeasy Mini Kit following the manufacturer’s instructions (Qiagen, Hilden, Germany). RNA quantification was done with a NanoDrop2000 spectrophotometer (Thermo Scientific, Waltham, MA, USA), and quality was assessed using the 2100 Bioanalyser (Agilent, Santa Clara, CA, USA). Only high-quality samples with a RIN value greater than or equal to eight (intact rRNA and no detectable trace of gDNA) were kept for subsequent steps. Final RNA concentration was measured with Quant-iT RiboGreen RNA Assay Kit (Invitrogen, Life Technologies, Carlsbad, CA, USA) before library preparation.

Individual libraries were prepared from 2µg of RNA using the TruSeq RNA sample preparation kit V2 (Illumina, San Diego, CA, USA) following the manufacturer’s instructions. Library size and concentration were evaluated using DNA High Sensitivity chip on the 2100 Bioanalyzer (Agilent, Santa Clara, CA, USA). Single read sequencing (100bp) was performed on the Illumina HiSeq2000 platform for the 72 libraries randomly distributed on a total of nine lanes (eight libraries/lane) at the McGill University and Genome Quebec Innovation Centre (Montreal, Canada).

### De novo transcriptome assembly and annotation

Raw sequencing reads were cleaned to remove adaptor and individual tag sequences, and trimmed using Trimmomatic v0.36 (Bolger et al. 2014). We applied a quality score threshold of 30 across a 10bp sliding window and removed all reads < 60 nucleotides in length after quality processing. Reads were merged among all individuals using FLASh v1.2.11 with default parameters (Magoc & Salzberg 2011), and used to assemble a *de novo* reference transcriptome using Trinity v2.2.0 suite (Haas et al. 2013). We aimed to build a combined orthologous gene composite reference (hereafter: common reference) for both the North American and European lineages in order to identify and compare orthologous genes involved in the divergence process between limnetic and benthic species across lineages. Successive filtering steps were applied to the *de novo* common reference transcriptome, contigs lacking an ORF longer than 200bp were discarded as well as redundant transcripts per ORF in favour of a unique ORF per transcript using TransDecoder v3.0.1 pipeline (-Transdecoder.LongOrfs) (TransDecoder 2016). In the absence of a reference genome and for comparative purposes with other salmonids transcriptomes, only the longest isoform per transcript was kept (Pasquier et al. 2016; Carruthers et al. 2018). Finally, a scaling factor of one Transcript per Million (TPM) was applied to normalize the raw reads count per gene for the gene expression analysis. We finally used a BlastX approach against the Swissprot database (http://www.uniprot.org) and the Ensembl *Danio rerio* database (Zv9) to annotate the filtered common reference. In parallel, we also assembled two separate lineage-specific transcriptomes for Lake and European whitefish following the same procedure as detailed above. Using normalized transcripts, we considered the reciprocal best hits within each transcriptome, for both *C. clupeaformis* and *C. lavaretus* to identify and discard paralogous genes (Carruthers et al. 2018). We then blasted each lineage-specific transcriptome to the common reference and discarded unmapped contigs, 98.7% and 98.2% of such orthologous hits for *C. lavaretus* and *C. clupeaformis*, respectively, thus avoiding imbalanced mapping between limnetic and benthic species of both lineages. We kept only common transcripts (*i.e.*, found in both lineages) that we refer to as orthologous genes in the common reference.

### Differential gene expression analysis

Individual reads were mapped to the orthologous gene common reference with Bowtie2 v2.1.0 (Langmead & Salzberg 2012) using the *-end-to-end* mode and reported multiple alignments were discarded. Then resulting Bam files were parsed to estimate individual reads counts with eXpress v1.5.1 (Roberts & Pachter 2012). Differential expression analysis were conducted with the R packages DESeq2 v1.14.1 (Love et al. 2014). In order to take into account the hierarchical structure of the studied populations, generalized linear models (glm) were built to allow for comparisons between benthic and limnetic species while integrating progressively lakes, regions and continents (hereafter called ‘phylogeographic’) effects, as covariates on gene expression. The final model for limnetic and benthic comparisons across continents was composed of: ‘∼Species*Lake*Continent’ in order to integrate interactions between ‘Continents’ and ‘Species’ in ‘Lake’, and interactions between ‘Lake’ and ‘Species’ within both ‘Continents’ successively. We then controlled for the presence of DEGs associated with phylogeographic effects by removing DEGs associated with ‘Lake’ and ‘Continent’ factors (*i.e.*, including interaction and intersection) from the list of identified DEGs, in order to keep only DEGs associated with the ‘Species’ factor. Inference of differentially expressed genes (DEGs) relied on normalized counts matrix by the integration of the size factor per library to correct for heterogeneity in sample sequencing depth. DEGs were determined by controlling for false discovery rate (FDR) as implemented in DESeq2 (Benjamini-Hochberg correction) (Benjamini & Hochberg 1995), with a threshold of a FDR<0.05. Then, GO enrichment analysis was performed with GOATOOLS (Tang et al. 2015), based on Fisher’s exact test. For all tested lists of genes, GO enrichment was associated with FDR<0.05 (Benjamini & Hochberg 1995) and a minimum of three genes represented per category.

### SNP genotyping and sequence divergence

In order to document the extent of polymorphism within and divergence between *C. clupeaformis and C. lavaretus* and among divergent sympatric benthic and limnetic species-pairs, individual reads were mapped (71.41% overall alignment mean success rate) to the common reference transcriptome using *Bowtie2* v2.1.0 ‘end-to-end’ alignment (Langmead & Salzberg 2012). Resulting SAM files were converted to BAM files and sorted using Samtools v1.3 (Li et al. 2009) and duplicates were removed with the Picard-tools program v1.119 (http://broadinstitute.github.io/picard/). The physical mapping information of reads to the reference was used for calling SNPs with *Freebayes* v0.9.10-3-g47a713e (Garrison & Marth 2012). Variable sites were considered for a minimum coverage of three reads per individual in order to process a site and for a minimum of two reads per individual to consider an alternative allele. We used the *vcffilter* program from *vcflib* (https://github.com/ekg/vcflib) to process the Variant Call Format (VCF) file obtained from *Freebayes*, in order to specifically retain biallelic SNPs with a phred scaled quality score above 30, a genotype quality with a phred score higher than 20, and an allele depth balance between 0.30 and 0.70. Following these quality control steps, we filtered the resulting VCF file using *VCFtools* v0.1.12b (Danecek et al. 2011), in order to remove miscalled and low quality SNPs for subsequent population genomics analyses. For each of the 12 populations, we kept loci with less than 10% of missing genotypes and filtered for Hardy-Weinberg disequilibrium using a *p*-value exclusion threshold of 0.01. Finally, we merged the VCF files from all the 12 populations, resulting in a unique VCF file containing 20,911 SNPs passing all the filters in each population. Since we did not apply any minor allele frequency threshold within populations, the final VCF represents a non-ascertained dataset of genetic variation. Intra-population nucleotide diversity (π) was estimated within non-overlapping 100bp windows (due to transcriptome data specificities; *i.e.*, N50 of 1,672bp) with *VCFtools* v0.1.12b (Danecek et al. 2011) on the VCF file of shared variant and invariant sites. We then reported the mean π per gene as the *a posteriori* mean of all windows per gene. Finally, we estimated the between-species nucleotide diversity (Dxy) with a custom *perl* script, using non-overlapping 100bp windows. Absolute divergence (Dxy) was calculated as the fraction of nucleotide differences between two sequences taken from two different species (Nei 1987; Cruickshank & Hahn 2014). As for the nucleotide diversity, we estimated the mean Dxy per gene from all windows contained in each gene.

### Shared polymorphism and historical relationships among lineages and species

Historical relationships among all populations were inferred using *Treemix* v1.12 (Pickrell & Pritchard 2012) applied to the VCF file containing 20,911 polymorphic SNPs. This program uses the covariance structure of allele frequencies between all tested populations and a Gaussian approximation for genetic drift to build a maximum likelihood graph relating populations with an ancestral genetic pool. The number of migration edges was determined empirically to improve the fit to the inferred tree. Migration edges among and between species may either reflect gene flow, or the retention of shared ancestral polymorphism among geographically isolated populations.

We then quantified and compared the amount of shared polymorphism retained at the lake, region and continental hierarchical levels. More precisely, we defined the number of SNPs that were shared and polymorphic among all populations for each hierarchical level (*i.e.*, between sympatric species at the lake level, among all populations from Norway, Switzerland and Maine at region level, and among all populations across both continents at continent level). We also tested for the increased probability of limnetic-benthic DEGs relative to non-DEGs to display shared polymorphism, which could hint to a possible role for selection in maintaining variation at these genes. We defined the proportions of DEGs with shared polymorphism (DEG_SP_) relative to the total number of DEGs (DEG_T_), and of Non-DEGs with shared polymorphism (NDEG_SP_) relative to the total number of Non-DEGs (NDEG_T_). From these two proportions we realised a ratio test to compare the relative proportion of shared polymorphism (SPRT: Shared Polymorphism Ratio Test) in each category of genes (*i.e.*, DEGs and NDEGs):

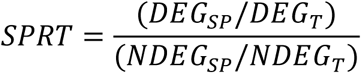

Confident intervals (CIs) of 95% were determined using 1000 bootstrapping iterations per comparison on the empirical dataset. Obtained ratios and associated CIs were compared to the expected ratio of one (*i.e.*, no difference in the amount of shared polymorphism between DEGs and NDEGs) to test for enrichment of shared polymorphism in DEGs (*i.e.*, SPRT>1).

### Detection of adaptive variation

The detection of adaptive variation using F_ST_-based approaches is challenging in study systems with complex population structures, which can be accounted for by multivariate outlier detection methods (Duforet-Frebourg et al. 2016; Luu et al. 2017). Redundancy Analysis (RDA) (Legendre & Legendre 1998) is an efficient constrained ordination method to detect (adaptive) variation under the effect of divergent selection, especially when the selection gradient is weakly correlated with population structure (Forester *et al.* 2018; Capblancq *et al.* 2018). In order to account for the hierarchical population structure, we used a conditioned (partial) Redundancy Analysis (cRDA), as implemented in the *vegan* v2.4-3 (Jari Oksanen et al. 2018) R package, to identify genes that diverge the most between limnetic and benthic species, independently of the population structure and hierarchical (region and continent) levels. Consequently, the first axis of the cRDA corresponds to the variance explained by the constrained ‘Species’ variable and the second axis corresponds to the PC1 of the PCA analysis (nested into the RDA) and represents the main axis of unconstrained variance. Because RDA can be sensitive to missing data, we only kept loci that contained no missing genotypes across populations, representing a total of 9,093 SNPs. Briefly, a RDA allows evaluating the variation that can be explained by the applied constraints. We conditioned the RDA to remove the effects of continents, regions and lakes to control for the hierarchical genetic structure. We tested the significance level of the cRDA with an analysis of variance (ANOVA), performed with 1,000 permutations. From the conditioned ordination, each SNP was assigned a locus score that corresponds to the coordinates used to ordinate points and vectors. Then, we identified outlier SNPs by putting significance thresholds at +/-2.6 and 3.0 standard deviations from the mean score of the constrained axis, corresponding to *p*-value thresholds of 0.01 and 0.001, respectively (Forester et al. 2018).

We also applied cRDA to gene expression data (*i.e.*, on the 32,725 orthologous genes from the common reference) and tested the significance of the constrained ordination model with an ANOVA using 1,000 permutations. We aimed at identifying co-varying DEGs between limnetic and benthic species after correcting for hierarchical population structure and applying a significance threshold on expression scores (*p*-value of 0.01 and of 0.001). This approach is expected to limit the local effect of highly expressed genes in the identification of outliers/DEGs if they were not associated to the constrained independent variables. Such co-varying DEGs could reflect the effect of polygenic selection acting in parallel between benthic and limnetic whitefish.

### Gene subnetworks analysis

In order to test for selection acting on sub-networks of genes involved in common biological pathways, we performed a gene network analysis designed specifically to detect polygenic selection (Gouy et al. 2017). The level of differential expression between limnetic and benthic whitefish captured by individual locus scores in the expression cRDA was scaled to a *z*-score, such that individual locus scores have a mean of 0 and a standard deviation of 1. We obtained Kegg Ontology (KO) for each transcript of the common reference with an Entrez gene ID annotation from the KASS (Kegg Automatic Annotation Server, http://www.genome.jp/tools/kaas/). Polygenic selection was tested using the R package *signet* (Gouy et al. 2017) on the *Danio rerio* and *Homo sapiens* KEGG databases (we present only the results obtained from the human database because of lack of power using the smaller *D. rerio* database). This package defines sub-networks of genes that interact with each other and present similar patterns attributed to selection, such as co-variation in expression level for genes involved in the same biological pathway. A null distribution of sub-network scores was generated by random sampling to create 10,000 sub-networks of variable sizes. Each pathway of the KEGG database was parsed to identify gene sub-networks with a high score using 10,000 iterations of simulated annealing. Finally, the *p*-value of the sub-networks showing parallelism (*i.e.*, for which we found evidence of differential expression between most to all limnetic and benthic populations) in gene expression was inferred based on the distribution of 10,000 permuted scores from the randomly generated sub-networks. We then tested for similarities among limnetic species for differential gene expression against benthic species across continents (sub-networks *P*<0.05).

### eQTL analysis

We related differential gene expression with sequence divergence to identify eQTLs. We thus generated a new VCF file containing shared loci among all the populations that showed polymorphism across continents (*i.e.*, trans-continental polymorphisms shared between *C. clupeaformis* and *C. lavaretus*), which corresponded to 2,240 SNPs. We extracted the 1,272 associated annotated genes and their expression level and tested for correlations between genotype and expression level (eQTL), using different models. We applied a linear model testing for gene expression variation in response to genotype variation by controlling for the lake and continent covariates as environmental effects to correct for population genetic structure (expression∼genotype+covariates). We compared the tested linear model to a theoretical simulated dataset, as suggested by (Shabalin 2012). This analysis was run using the R package *MatrixEQTL v2.1.1* (Shabalin 2012). We identified significant (FDR<0.05) *cis-*eQTL by focusing on SNPs affecting the expression level of the gene to which they were physically linked, from the output of the linear model including all covariates.

## RESULTS

### Reference transcriptome assembly

A total of 1.74 × 10^9^ 100bp single-end reads (average of 24.22 × 10^6^ reads per individual) were generated from 72 individuals (Table S1). Filtered libraries (1.69 × 10^9^ reads) were used to *de novo* assemble a composite reference transcriptome. Starting from a raw assembly of 277,194 contigs (mean length = 768bp; N50 = 1,347bp), we kept only the longest ORF per transcript, which reduced the initial assembly by 66.9% (91,715 contigs remaining, see Table S2). Keeping the longest isoform per transcript and normalizing the reads distribution resulted in a composite reference transcriptome composed of 54,514 contigs (mean length = 1,121bp; N50 = 1,672bp) with 79% of uniquely annotated transcripts (43,501 contigs). This liver-specific assembly is consistent with the number of transcripts assembled using several organs separately in one *C. clupeaformis* male and one *C. lavaretus* female (range: 66,996 - 74,701, respectively (Pasquier et al. 2016)). Comparing transcriptomes separately assembled in *C. clupeaformis* (55,104 contigs, with 83.8% uniquely annotated) and *C. lavaretus* (58,321 contigs, with 73.9% uniquely annotated) to identify orthologous genes and filtering out paralogous genes (*i.e.*, self-mapped transcripts hits, 15%), we ended up with a common reference transcriptome of 32,725 annotated contigs (comparable to (Carruthers et al. 2018)) that were used for downstream analyses of gene expression and sequence divergence. The common reference transcriptome N50 was 1,797bp with a contig size distribution ranging from 297bp to 13,427bp and a mean contig size of 1,185bp.

### Genetic relationships among populations

Characterizing the genetic relationships among the studied populations with TreeMix indicated the presence of shared polymorphisms maintained across the entire hierarchical genetic structure (Fig 1 and Fig S1). The different hierarchical levels were composed by limnetic-benthic species-pairs (hereafter, Species-pair level) in both North America and Europe (the lake-level was removed for more clarity in this analysis). Similarities in branch length at the Species-pair level reflected the similar degrees of genetic differentiation among species-pairs from different regions consistent with a relatively similar timing of divergence and postglacial admixture among species-pairs from both continents. The second hierarchical level was composed by intra-continent regional divergence (hereafter, Region level), which was represented by the two European regions; Central alpine (Switzerland) and Fennoscandinavia (Norway). The highest hierarchical level of divergence was between the two sister-lineages *C. clupeaformis* and *C. lavaretus*. The sharing of ancestral polymorphism between geographically isolated taxa across continents was captured by the inferred migration link connecting two limnetic populations from Maine and Norway (Fig 1). This link indicates an excess of shared ancestral polymorphism after accounting for drift along the population tree, which could indicate the presence of balanced polymorphisms across continents or past admixture events.

### Trans-continental polymorphism quantification

Given the evidence for shared polymorphisms maintained among species from different continents, we documented the overall extent of trans-continental polymorphism. Trans-continental polymorphism corresponds to ancestral variation shared among all populations of limnetic and benthic species from North America (*C. clupeaformis*) and Europe (*C. lavaretus*), that is, loci that are polymorphic in all populations on both continents. Among the 20,911 SNPs initially obtained after genotyping and filtering steps, we identified 2,241 SNPs (10.7%) distributed among 1,251 genes (3.8%) that met our criteria of trans-continental shared polymorphic loci. The genes containing trans-continental polymorphisms showed a significantly higher mean level of nucleotide diversity (π) per gene within species (Wilcoxon signed-rank test, *P*<0.001, Fig S2), and a higher mean level of absolute sequence divergence (Dxy, Nei 1987) per gene between limnetic and benthic species (Wilcoxon signed-rank test, *P*<0.001, Fig S3) compared to genes with no trans-continental SNPs. These results point to the influence of evolutionary processes (after controlling for artefact from the data, Fig S4) acting differentially between these two categories of genes.

### Parallel genetic differentiation

We used conditioned ordination to test whether divergence at the limnetic/benthic species-pair level involves parallel changes in allele frequency across sister-lineages from different continents (*C. clupeaformis*-*C. lavaretus*). We conditioned the ordination to account for the hierarchical genetic structure among populations (Lake, Region and Continent). The cRDA thus allowed the identification of variants associated with limnetic-benthic species divergence, explaining 2.8% of the total genetic variance (ANOVA, *F*=1.259, *P*=0.001), after controlling for the variance explained by regional and continental population structure (Fig 2A and Fig S5). The distribution of individual locus scores on the first cRDA axis discriminating all limnetic and benthic samples allowed identifying 348 outlier markers (*P*<0.001, 3.0 s.d.) showing parallel allele frequency differences between limnetic and benthic species across both continents (Fig S6). These 348 SNPs, which represent 15% of the 2,241 of the trans-continental polymorphic loci may be interpreted as being enriched for shared genetic bases of limnetic-benthic species divergence across continents. Gene ontology (GO) analysis of transcripts associated with parallel outlier SNPs revealed significant enrichment (*P*<0.001) in metabolic process (*i.e.*, catabolism), immune system process and developmental process, among others (TableS1).

**Fig 2.**
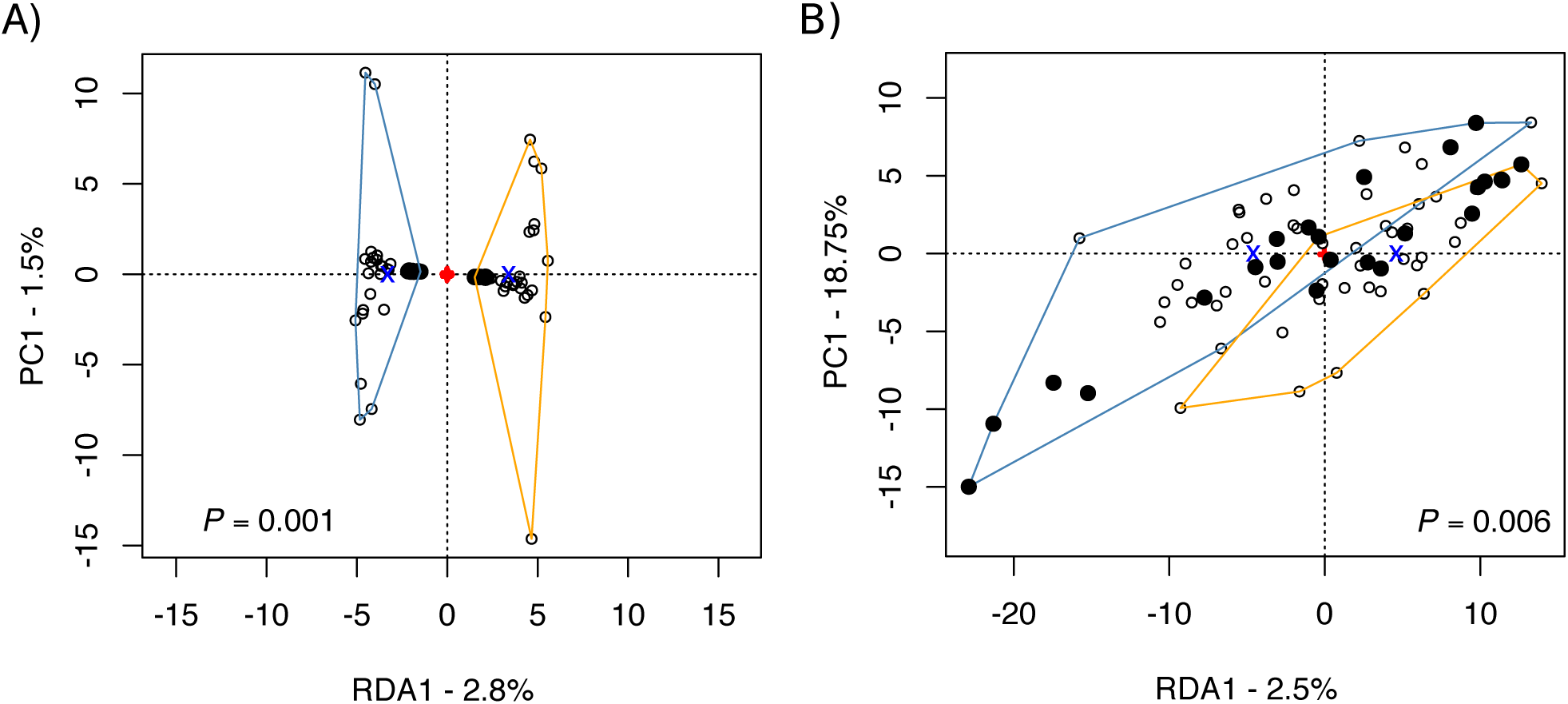
Conditioned redundancy analysis (cRDA) clustering individuals per species. (all limnetic vs. all benthic from both continents). The cRDA was used to capture divergence between limnetic and benthic species (along RDA axis 1) while correcting for three hierarchical levels of population structure (lake, region and continent) based on A) genotypes and B) gene expression levels. Orange and blue clusters correspond to benthic and limnetic species, respectively. Circles represent individuals. Open and filled circles represent the *C. lavaretus* and *C. clupeaformis* lineages, respectively.

### Differential gene expression between Limnetic and Benthic Species

Using the benthic whitefish populations as the reference level, we quantified differentially expressed genes (DEG) between limnetic and benthic whitefish at the Lake and Region levels using multivariate glm. Despite different sampling locations, storage methods and sequencing runs, we did not observe any sign of experimental bias in our transcriptomic data (Fig S7). Our expectation was a decreasing number of shared DEGs in higher comparisons levels, due to a reduced fraction of shared regulatory variants. In North America, Cliff and Indian Lakes, showed 3,175 (9.7%) and 238 (0.7%) significantly differentially expressed genes (False discovery rate; FDR<0.05) between sympatric limnetic and benthic species, respectively (Fig 3). In both lakes, approximately twice as many genes showed higher expression in the limnetic species compared to the benthic species (Cliff Lake: 2,001 *vs*. 1,174, *χ*^*2*^ test, *P*<0.001; Indian Lake: 159 vs. 79, χ2 test, P<0.001) (Fig S8). The lower level of DEGs identified in Indian Lake was likely associated to lower level of differentiation between species-pairs, but also to a much higher inter-individual variance in this lake (Fig S8). While we found 44 common DEGs, using a less stringent significance threshold (*q*-value<0.1 instead of 0.05) led to the identification of 1,926 DEGs in Indian Lake, 318 of which were shared with the 3,175 DEGs from Cliff Lake. In Langfjordvatn and Skukkebukta lakes from Norway, 276 and 112 significant DEGs were identified between limnetic and benthic whitefish, respectively. Contrary to North America, more genes showed lower expression in limnetic populations (45 *vs.* 231, *χ*^*2*^ test, *P*<0.001 in Langfjordvatn; 44 *vs*. 68, *χ*^*2*^ test, *P*=0.023, in Skrukkebukta) (see also Fig. S8). In Swiss lakes, 3,727 and 1,392 genes showed a significantly different expression level between limnetic and benthic species in Lake Lucerne and Lake Zurich, respectively. In contrast with North America and Norway however, a similar number of genes showed lower- and higher-expression in the limnetic species compared to benthic species in both lakes (1,870 *vs.* 1,857, *χ*^*2*^ test, *P*=0.831 in Lake Lucerne and 691 *vs*. 701, *χ*^*2*^ test, *P*=0.787, in Lake Zurich).

**Fig 3.**
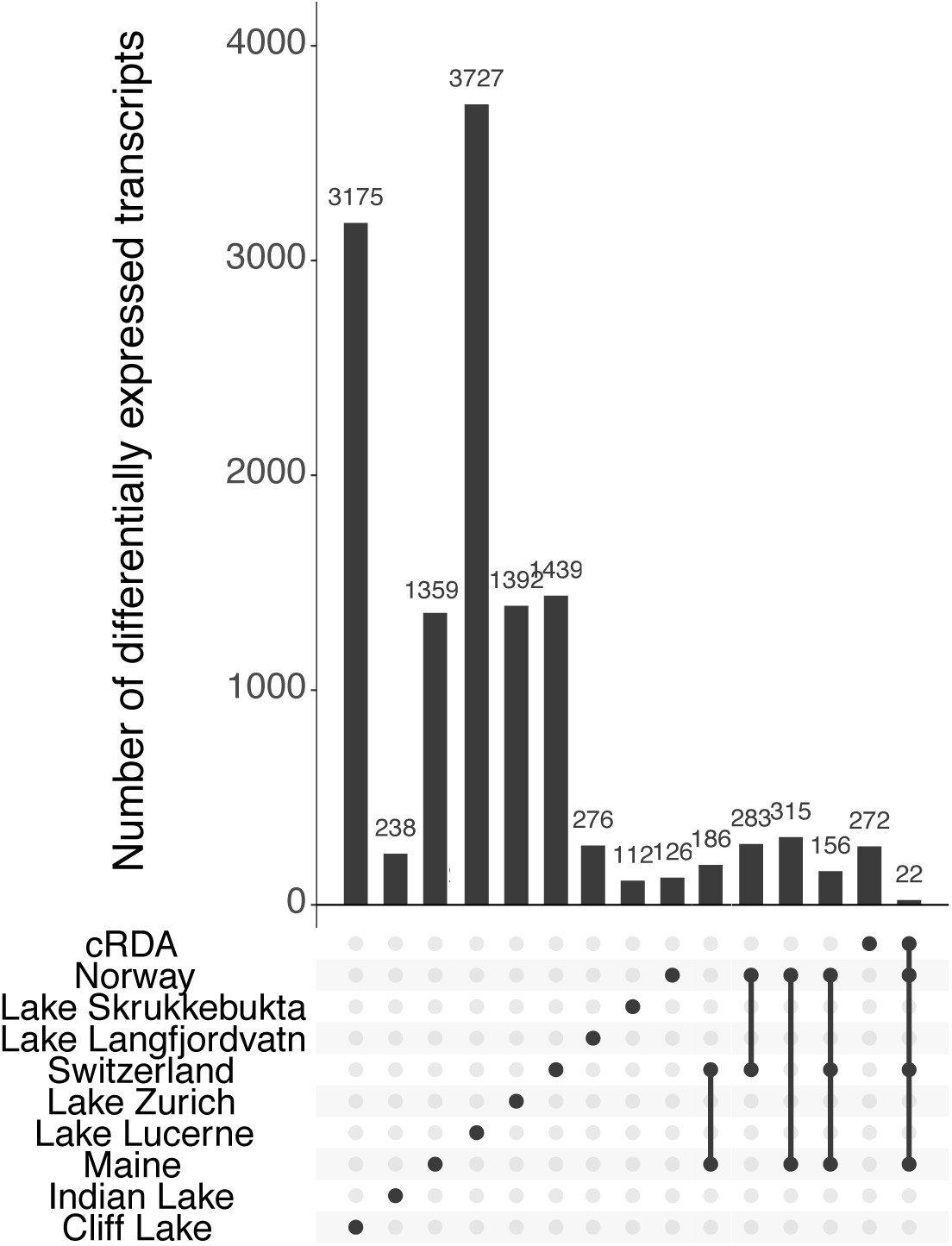
Frequency of shared differentially expressed transcripts between species across hierarchical levels. Limnetic/Benthic comparisons are indicated by the dots (intra-lake and intra-region). Linked dots represent the combinations of pooled populations per species for inter-species comparison levels (inter-regions and inter-continents). The number of significant differentially expressed genes (DEGs) associated to species divergence (FDR<0.05) per comparison is indicated on top of each bar, as determined from the multivariate tests, from the cRDA for all limnetic to all benthic comparison, as well as the overlap of DEGs between cRDA and DESeq2 (last column).

To document the extant of parallelism at the Region level (*i.e.*, common DEGs between species across lakes within a given region), we considered the proportion of DEGs between limnetic and benthic whitefish while integrating the lakes as covariates in a multivariate glm. Thus, the degree of parallelism in DEG was most pronounced in Swiss lakes whereby 1,439 parallel DEGs were found between limnetic and benthic species among lakes. These parallel genes showed a higher proportion of down-regulated genes in limnetic species (625 vs. 814, χ2 test, P<0.001) (Fig S9). Similarly in Maine, 126 parallel DEGs were identified between limnetic and benthic species from Indian and Cliff lakes. These genes showed a higher gene expression in the limnetic species than in the benthic species (818 vs. 541, χ2 test, P<0.001) (Fig S9). Finally, 126 parallel DEGs were identified between limnetic and benthic whitefish from the two Norwegian lakes. Here, however, parallel DEGs comprised a significantly higher proportion of down-regulated genes in the limnetic species (53 vs. 239, χ2 test, P<0.001) (Fig S10).

GO enrichment analysis at the Lake and Region levels provided evidence for parallelism in biological functions. Indeed, DEGs were significantly enriched (FDR<0.05, see Table S3) in limnetic species for immune system response, detoxification and antioxidant activity in both North America and Europe. Moreover, we found enrichment in genes associated to growth and development at both Lake (Indian, Lucerne, Skrukkebukta and Langfjordvatn lakes) and Region levels (Maine) in benthic whitefish. DEGs were also enriched in limnetic species for metabolic processes, electron carrier activity and catabolic processes, which are associated with differences in the metabolic rate between limnetic and benthic species (Laporte *et al.* 2016; Dalziel *et al.* 2017b; a).

At the Continent level (integrating Region and Continent as covariates), we found 156 parallel DEGs between limnetic and benthic species between both continents. Again, these 156 genes showed similar proportions of up- and down-regulated genes in all limnetic whitefish compared to all benthic whitefish (72 *vs*. 84, *χ*^*2*^ test, *P*<0.299; Fig S10). From enriched biological functions included metabolic process (*P*=0.016) and antigen binding (*P*=0.021) associated with immune response (*e.g.*, Immunoglobulin domain) and cellular metabolic process (*P*=0.002) (*e.g.*, SPRY-associated domain). DEGs were also enriched for oxido-reductase activity (*e.g., TSTA3*, a gene able to activate fructose and mannose metabolism via oxydo-reductase step, involved in Glycolysis; *Hsp90*, a gene responding to environmental stress with effects on growth).

### Enrichment in trans-continental polymorphism in DEGs

The excess of ancestral polymorphism shared among limnetic and benthic species across both continents suggests the existence of a mechanism responsible for the maintenance of balanced ancestral variation, against the stochastic effect of drift in each lineage. In order to further test if the retention of ancestral polymorphism could be linked to differential selection on adaptive traits between limnetic and benthic species, we tested if *cis*-regulating regions (*i.e.*, regions physically linked to transcripts) of DEGs show an increased probability of having shared polymorphisms. The shared polymorphism ratio test (SPRT), which compares the proportions of shared polymorphism in DEGs to non-differentially expressed genes (NDEGs), revealed an enrichment of shared polymorphism in DEGs at the three hierarchical levels (Fig 4), and a parallel analysis suggested that DEGs do not have a higher level of expression (*i.e.*, a higher sequencing depth) than NDEGs (Fig S11).

**Fig 4.**
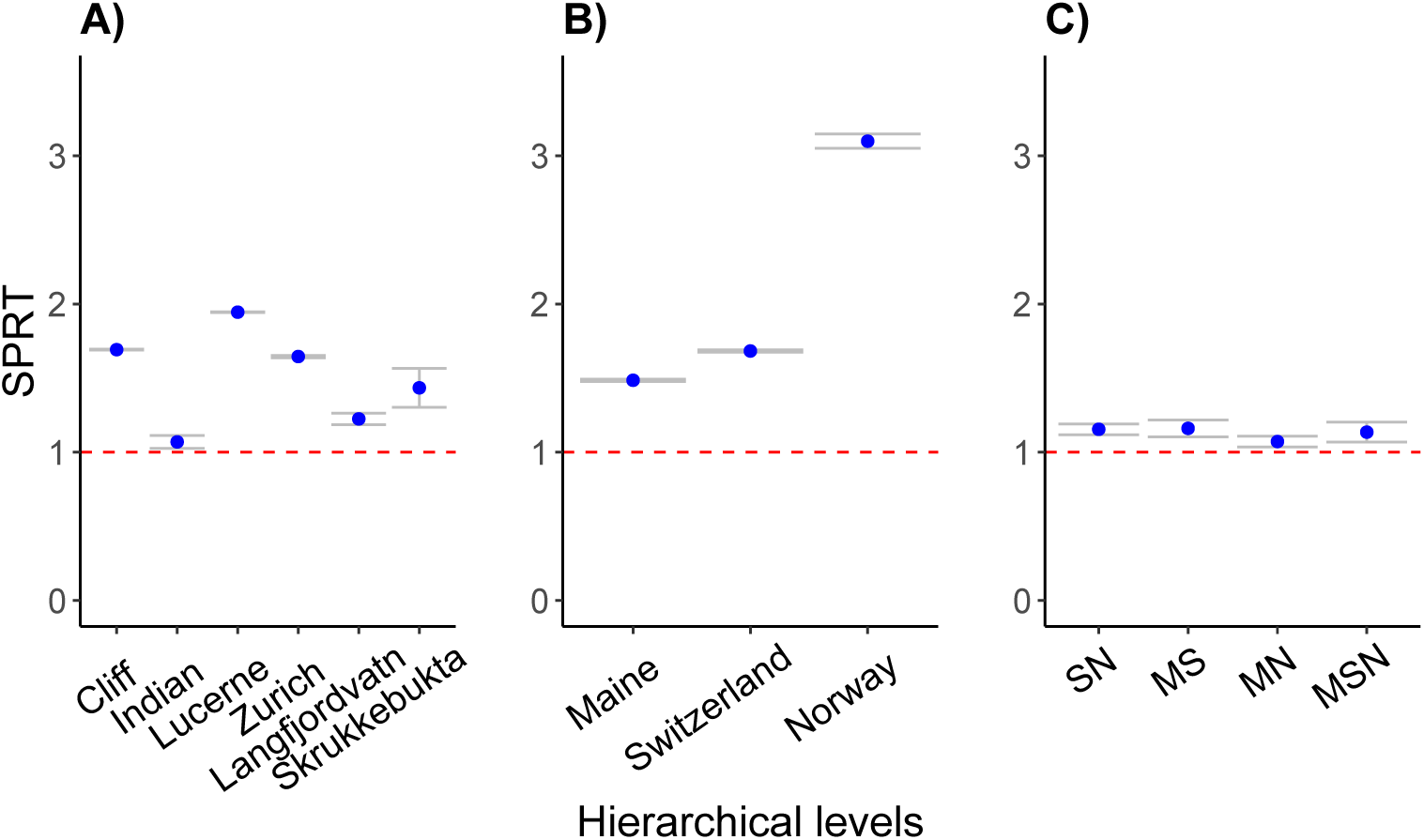
Shared polymorphism enrichment in DEGs between species compared to Non-DEGs. Test ratio (SPRT) of the proportion of shared trans-continental polymorphism in DEGs compared to non-DEGs, for A) intra-lake, B) inter-lake within regions and C) inter-regions (SN: Switzerland/Norway, MS: Maine/Switzerland, MN: Maine/Norway and MSN: Maine/Switzerland/Norway) comparisons. All ratios are above the red-dashed line (y=1), which illustrates the general enrichment of shared polymorphism in DEGs compared to non-DEGs. Grey bars indicate the 95% confidence interval associated with the observed value obtained using 1000 bootstrap resampling of the dataset.

### Identification of gene sub-networks showing patterns of parallel gene expression

A credo performed on the expression data of the 32,725 orthologous genes revealed that 2.5% of total variance in expression was explained by net differences between limnetic and benthic species across continents (ANOVA, *F*=2.516, *P*=0.006, Fig 2B), while accounting for the hierarchical population structure. The z-scored distribution of the gene expression on the first RDA axis constrained for divergence between limnetic and benthic whitefish ranged from [-4.14; 3.99] (Fig S12). Applying two different significance thresholds (*P*<0.01 and *P*<0.001) allowed identifying 272 (*P*<0.01) and 66 (*P*<0.001) putative outliers DEGs. These were significantly enriched for biological regulation (*P*<0.001) and metabolic process (*P*<0.001) in both sets of genes, and growth (*P*=0.026) for the subset of 272 genes (Table S3). Twenty-two out of the 156 (14%) parallel DEGs at the Continent level (which is more than expected by chance, hypergeometric test, *P*<0.001) identified with the *glm* analysis overlapped with the 272 DEGs from the cRDA on gene expression, including the previously mentioned genes *TSTA3* and *Hsp90*.

We then investigated the polygenic basis of transcriptomic differences using genes expression scores defined by the ordination analysis. The *z*-scored transformation of cRDA’s gene expression scores was used as a quantitative measure for assessing the extent of parallel expression between limnetic and benthic whitefish across continents. A total of 22,188 out of the 32,725 orthologous genes (from the common reference) that were successfully annotated with an Entrez gene ID were analysed with *signet* based on information from KEGG databases. In *signet*, gene sub-networks (*i.e.*, genes showing patterns of convergence within limnetic populations) were identified for each pathway and we considered the significance of sub-networks (*P*<0.05) in the analysis against the *Homo sapiens* KEGG database.

Ten metabolic pathways with significant sub-networks of genes were identified (Table S4). Five of these pathways shared genes and were therefore merged together to identify genes showing convergent patterns among the significant sub-networks (Fig 5, Fig S13), mainly for peripheral genes (Fig S14). From the ten identified metabolic pathways composed of 73 parallel DEGs between species, three categories of metabolic functions were represented. The first category corresponded to energetic metabolism (*e.g.*, pentose phosphate pathway, glycerolipid metabolism, nicotinate and nicotinamide metabolism, FoxO signalling pathways) which is involved in regulation of glycolysis and energy production (from ATP to NADH). The second category was the detoxification metabolism and immune system (*e.g.*, CYP450 and glutathione metabolism), which is mainly associated to detoxification and oxidative stress, maintaining cell integrity by preventing damage due to reactive oxygen species (ROS). The third category was the cell cycle metabolism and control (*e.g.*, FoxO signaling pathways, purine and pyrimidine metabolism, cAMP signalling pathway, PI3K-Akt signaling pathway). These pathways play critical roles in regulating diverse cellular functions including metabolism, growth, proliferation, survival, transcription and protein synthesis.

**Fig 5.**
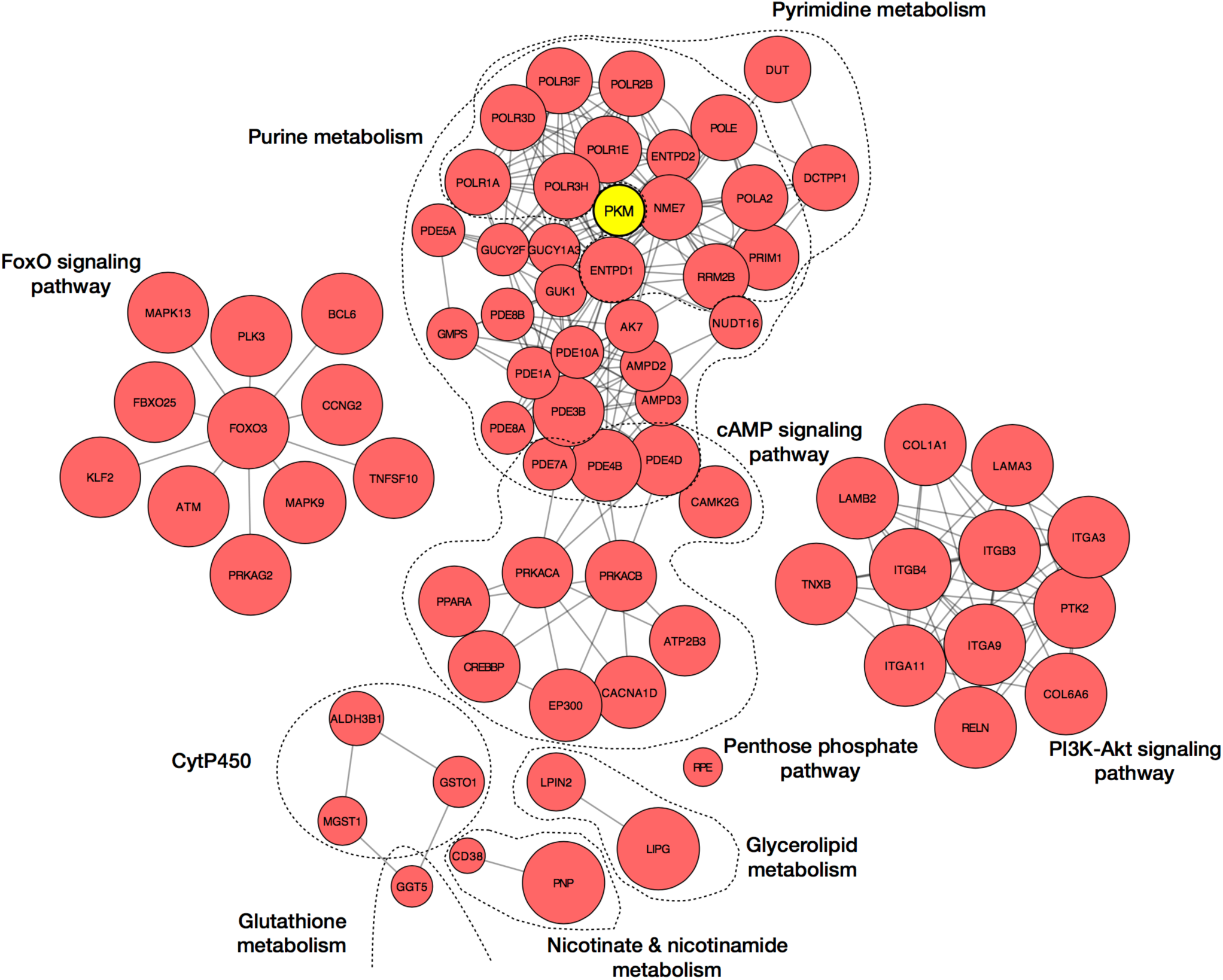
Merged significant subnetworks for Limnetic/Benthic species comparisons. Detailed subset of differentially expressed genes between species showing convergence across the studied system, separately displayed for each metabolic pathway. Pyruvate kinase (PKM) gene expression (in yellow) is associated with a *cis*-eQTL in its 3’UTR. Gene annotation is based on the Ensembl nomenclature. For each node represented by a gene, the relative size is proportional to the contribution score of the associated gene to the significance of the metabolic pathway. The score for each gene corresponds to a probability of convergent adaptation between individuals of the same species.

### cis-eQTL markers associated with expression

We finally tested whether the level of expression was associated with SNPs located within a given gene. We found 451 SNPs significantly (P < 0.05) associated with the level of expression of the gene (*i.e.*, UTR and coding sequence) to which they belong (*cis-*eQTL). That is, the level of expression significantly varied among the three possible genotypes at a given SNP. Controlling for multiple tests and testing for deviation in the distribution of p-values due to unaccounted covariates (Fig S15), we retained 134 significant (FDR<0.05) *cis-*eQTL across continents associated to differences between limnetic and benthic species.

We identified SNPs and genes showing overlap between the different analyses. Thus, two significant *cis-*eQTL overlapped the 20 outliers SNPs from the cRDA based on genetic variation (hypergeometric test, *P*<0.001). They were physically linked to the complement factor H (*CFH*, Entrez 3075; Fig 6B) and Protein Kinase AMP-Activated Non-Catalytic Subunit Beta 1 (*Prkab1*, Entrez 19079; Fig 6C). These genes are respectively involved in immune response and in regulation of the cellular energy metabolism. Moreover, among the 134 significant *cis-*eQTL, 19 genes were shared with genes identified in the polygenic subnetwork analysis (hypergeometric test, *P*=0.006). However, only the pyruvate kinase gene (*PKM*, isoform M2; Entrez 5315) remained significant in both (sub-networks and eQTL) analyses (Fig 4). This gene encodes a protein involved in glycolysis, which generates ATP and pyruvate. The level of expression of this gene was higher in heterozygous and homozygous individuals for the minor allele (Fig 6A and Fig S16, linear model, *P=*0.001). Finally, we inferred the gene structure (*i.e.*, identification of 5’ and 3’ UTR, exonic and intronic regions) of our *de novo* assembled transcriptome and more particularly the *PKM* gene. We localised the variant affecting the level of expression of the *PKM* gene in its 3’UTR region, which could impact the regulation of the transcription of this gene.

**Fig 6.**
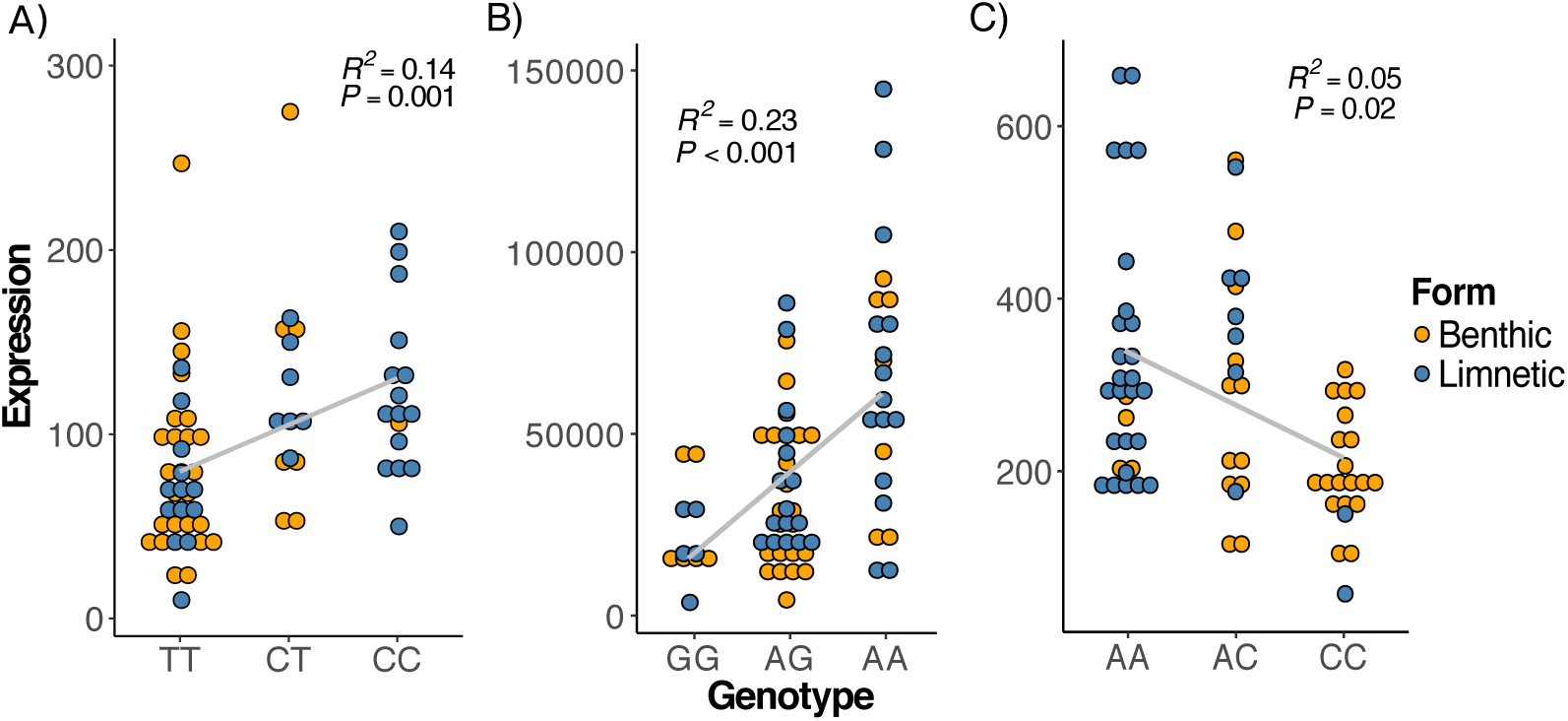
Associations between significant *cis*-eQTL genotypes and the level of expression (relative counts) of three genes in limnetic and benthic whitefish species, independently of their geographic origin. Three examples of transcripts abundance per individual (circles) varying with genotypes for a *cis*-eQTL located in 3’UTR of A) pyruvate kinase (PKM), B) complement factor H (CFH) and C) Protein Kinase AMP-Activated Non-Catalytic Subunit Beta 1 (Prkab1) genes. The grey line corresponds to the linear model fitted to the data and associated statistics (coefficient of determination: *R*^*2*^ and *p*-value: *P*) detailed in each panel. Individuals from benthic and limnetic sister-species are represented in orange and blue, respectively.

## DISCUSSION

*C. clupeaformis* (North America) and *C. lavaretus* (Eurasia) sister lineages have been geographically isolated for the past 500,000 years (Bernatchez & Dodson 1991; 1994; Jacobsen *et al.* 2012). Yet, they maintained similar habitat preferences in cold freshwater lakes (Bernatchez & Dodson 1991; Douglas *et al.* 1999; Østbye *et al.* 2005; 2006), with a frequent occurrence of sympatric species pairs being respectively associated to benthic and limnetic ecological and trophic niches (Lu & Bernatchez 1999; Amundsen *et al.* 2004; Kahilainen & Østbye 2006; Landry *et al.* 2007; Häkli *et al.* 2018). In both *C. clupeaformis* and *C. lavaretus*, historical demographic events and selective processes initiated species diversification (Rougeux *et al.* 2017; 2019), and resulted in a repeated ecological specialisation to limnetic and benthic habitats in each region. Here, the analysis of gene sequence divergence and differential expression in limnetic-benthic species has the potential to provide new insights into the genomic bases of parallel adaptation and parallel ecological speciation.

A salient result from our analysis is that pairs of limnetic and benthic species from independent divergence events exhibit parallelism in DEGs associated with repeated divergent adaptation to different ecological niches (*i.e.*, limnetic and benthic niches). The identification of 156 significant parallel DEGs involved in energetic metabolism, immune response, cell cycle and growth is congruent with previous transcriptomic analysis conducted in *C. clupeaformis* on the same organ tissue but with low resolution methods (microarrays with a reduced representation of transcripts), highlighting life history trade-offs between growth and energetic costs associated with occupying the limnetic niche (St-Cyr et al. 2008; Jeukens et al. 2009). Those congruent results also highlight the fact that despite differential storage methods, our dataset was not affected by any of the controlled batch effects and that such putative source of stochasticity (increased variance) did not significantly affect our inference of differential expression and parallelism between species. Moreover, our results show that seeking to detect parallel DEGs based on a single gene approach may lack the power to detect polygenic changes in expression levels, as it could be expected for the complex phenotypic traits involved in the divergence of these species-pairs. Indeed, a gene-by-gene approach may be too conservative and not well adapted to capture subtle parallel expression differences at genes involved in the same biological pathways under selection. The statistical approach employed to detect polygenic selection based on transcript abundance co-variation allowed us to integrate this level of information. Consistent with results from the negative binomial glm, the RDA allowed identifying a parallel genetic basis of phenotypic and ecological divergence by revealing parallel DEGs between limnetic and benthic species. Moreover, we found that these DEGs are involved in several metabolic pathways belonging to energetic, growth, cell cycle metabolisms and transcription factor, regulating genes associated with energetic metabolism, as observed in Jacobs *et al.* (2018). This approach also allowed detecting congruent expression signals at the integrated pathway scale, where the same effect on the selected phenotype can be achieved via regulation of different genes, because of the complexity and redundancy of the multi-genic regulatory systems (Yeaman 2015). The accumulated results coupled with previous analyses on this system thus highlight the repeated action of natural selection on gene expression patterns (St-Cyr *et al.* 2008; Jeukens *et al.* 2009; Jeukens & Bernatchez 2011), as well as on partially shared polygenic bases of phenotypic traits (Gagnaire *et al.* 2013a; b).

In addition to trans-continental parallelism in interspecific divergence of transcript abundance, we found parallel differentiation between limnetic and benthic species also at the gene sequence level. Indeed, from identified outliers between replicate species-pairs, 15% were defined as parallel outliers (*i.e.*, shared among all species comparisons), consistent with what has been found in other systems (ranging from 6% to 28%; (Deagle et al. 2012; Westram et al. 2014; Ravinet et al. 2015; Le Moan et al. 2016; Meier et al. 2018)). We also identified 1,251 genes exhibiting shared polymorphism across the inter-continental complex of *Coregonus* lineages. Similar patterns were observed in the tree species *Populus tremula*, in which genes in the networks’ peripheries (*i.e.*, with lower level of connectivity) were more likely to be enriched in polymorphism due to reduced evolutionary constraints (Mähler et al. 2017). For these genes, patterns of genetic diversity and DEGs enrichment between limnetic and benthic species suggest the action of divergent selection in the presence of gene flow (Charlesworth et al. 1997) globally maintaining alleles associated with different expression levels between sympatric species. The fact that absolute genetic divergence between sympatric limnetic and benthic species in genes with trans-continental shared polymorphism was elevated compared to other genes may reflect the active maintenance of these alleles over the long term (*i.e.*, since the lineages divergence). Such long term maintenance could result from the interaction of divergent selection and admixture between sympatric species-pairs (Han *et al.* 2017; Ma *et al.* 2017). The action of gene flow between sympatric limnetic and benthic whitefish may indeed further contribute to maintain the alleles favoured in each species in a balanced state within each lake, thus protecting polymorphism from being lost at a global scale even across continents. This could be eased by the apparently highly polygenic nature of the traits under divergent selection, meaning that the intensity of selection acting on each underlying locus could be weak (Le Corre & Kremer 2012). Therefore, even a modest amount of gene flow could possibly maintain a balanced polymorphism within each species-pair. This, however, remains to be investigated more formally, since strong divergent selection without migration within lakes would also help the maintenance of polymorphism across continents. Moreover, such genomic patterns could be directly caused by the sorting of sieved ancestral balanced polymorphism, which genomic signatures would mimic patterns built by other evolutionary process (*e.g.*, gene flow, as discussed above) (Guerrero & Hahn 2017).

The identification of orthologous genes with trans-continental polymorphism associated with differential expression between benthic and limnetic species supports the existence of *cis*-acting SNPs on transcripts abundance. Moreover, the characterisation of DEGs enriched in shared polymorphism across continents suggests the long-term action of some form of balancing selection, maintaining ancestral polymorphisms that predate the onset of regional and continental divergence of the different limnetic-benthic species-pairs. Consistent with theory and empirical studies (Zheng et al. 2011), our analysis of orthologous genes supports a role of polymorphism originating from standing genetic variation both in protein coding sequences (CDS) and regulatory motives (*e.g.*, untranslated regions UTRs) in the process of adaptive divergence between limnetic-benthic whitefish sister species (Zheng et al. 2011). For instance, we found a parallel *cis*-eQTL in the 3’UTR of the pyruvate kinase gene (*PKM*), affecting the relative expression level of this gene between species. The *PKM* isoform M2 corresponds to a glycolytic enzyme (iso-enzyme) that is expressed in liver tissue. Given the importance of 3’UTRs in regulating the transcription process and transcripts abundance (Merritt *et al.* 2008; Wittkopp & Kalay 2011), this 3’UTR SNP could be under divergent selection between limnetic and benthic species and therefore protected from being lost by drift within populations over the long term within any given limnetic-benthic pair as hypothesized above. Consequently, it is likely that such a *cis*-eQTL could have been recruited from standing genetic variation by natural selection, increasing in frequency in limnetic whitefish on both continents, while modifying the level of expression of a central gene in energetic metabolism.

The inferred module of gene co-expression from liver tissue allowed quantifying a partial view of the gene co-expression network associated with species phenotypic differentiation. Interactions between nearby genes within the module could result in *cis*-regulation affecting the level of expression of other genes and ultimately, affect the activity of genes farther in the genome (Boyle et al. 2017). It is noteworthy that genes interactions between metabolic pathways are conserved among sister-lineages. Indeed, it has been shown that co-expression modules are maintained through evolutionary times despite variation in the set of regulatory genes that activate them (Tanay et al. 2005). Moreover, those patterns of gene expression changes, maintained across the system, could be associated to genes involved in local adaptation by which co-evolving traits are integrated into the same module (Wagner et al. 2007). Despite the partial portrait of the polygenic basis of phenotypic differentiation between species, we stress that no gene of main effect (*i.e.*, hub gene) was identified in the module. Such patterns would suggest that modularity in gene expression and genes interaction into a module can recruit less constrained genes without affecting highly constrained central genes (Wagner 1996), while peripheral genes could be directly and indirectly involved in the co-expression network via a ‘hub-gene’ effect in the initial metabolic pathways of the recruited gene. However, further investigations on quantifying the gene-gene interactions or protein-protein interactions (PPIs), in order to infer individual gene constraint levels or position in the module, should be realized in a more formal framework on several tissues. Thus, this could allow identifying a most complete picture (qualitatively and quantitatively) of the gene co-expression network associated with phenotypic differentiation between limnetic and benthic species.

In conclusion, our study provides a quantitative assessment of DEGs and gene sequence divergence based on an extensive transcriptomic dataset, enabling to infer the effects of polygenic divergent selection acting on complex traits that diverge between sympatric benthic and limnetic species, within both the *C. clupeaformis* and *C. lavaretus* species radiations (Gagnaire et al. 2013b; Laporte et al. 2015). Our results also extend previous findings by revealing patterns of parallelism between species on two continents, derived from two evolutionary lineages that diverged at least half a million years ago. Furthermore, they show the effects of polygenic selection on genes associated with fundamental and constrained metabolic pathways, such as functions associated with energetic metabolism (Dalziel *et al.* 2017a). Due to the additive effects of multiple genes in controlling the expression of polygenic phenotypic traits, the probability of identifying a shared genetic basis from standing genetic variation (likely to increase with the number of genes involved) is higher compared to the alternative *de novo* mutation to generate local polymorphism. This suggests an important contribution of ancestral polymorphism in the repeated evolution of sympatric species pairs. This was illustrated by the identification of a genetic variant in the UTR gene region associated with phenotypic differences between species, as previously reported in other taxa (Schluter *et al.* 2004; Wittkopp *et al.* 2004; Jones *et al.* 2012; Verta *et al.* 2016; Uebbing *et al.* 2016; Wang *et al.* 2017). The resolution of future studies could be enhanced using a comparative whole-genome resequencing approach to provide a more detailed understanding of the genomic architecture of phenotypic differences between species, and the role of old standing variants in ecological speciation.

## ACKNOWLEDGEMENTS

Jérémy Le Luyer, Ben J. G. Sutherland provided valuable advices and discussions on transcriptomic analysis, Martin Laporte for inspiring and stimulating discussions about the lake whitefish system, thanks guys. We thank Alexandre Gouy for support with *signet* in the early times of the package, as well as Kyle W. Wellband for commenting on an earlier version of the manuscript. We thank Shripathi Bhat and the Freshwater ecology group at UiT for participating with the sampling in Norway. Finally we are grateful to AE Shawn Narum and two anonymous referees for their constructive and helpful comments on an earlier version of the manuscript.

## AUTHOR CONTRIBUTIONS

CR and LB designed the project. KP and OS shared samples from Norway and Switzerland, respectively. CR produced and analyzed the data. CR drafted the manuscript and all authors contributed to the writing and approved the final draft of the manuscript.

## DATA ACCESSIBILITY

Raw sequence data are available through the NCBI sequence read archive (SRA) database under accession SRP136771. Scripts for analysis are available under the github repository: https://github.com/crougeux.

## Supporting Information Legends

**S1 Fig. Genetic clustering of limnetic and benthic species’ populations.** A) Principal component analysis (PCA) generated from 20,911 SNPs. Individuals from Cliff (C), Indian (I), Langfjordvatn (LF), Skrukkebukta (SK), Zurich (Zu) and Lucerne (Lu) lakes for limnetic (-D) and benthic (-N) species are projected along PC1 discriminated populations of *C. clupeaformis* form *C. lavaretus*, and PC2 lakes from Norway and Switzerland. B) Similarity tree decomposing the individual genetic distance across the system. Less differentiated species-pairs clustered together within region (*e.g.*, Norway in green and Indian Lake in Maine region, black).

**S2 Fig. Distribution of genetic diversity (∏) for two categories of genes.** Genes with trans-specific polymorphism (yellow) show higher genetic diversity among species than genes without trans-specific polymorphism (Student’s *t*-test, *p*-value<0.001).

**S3 Fig. Distribution of absolute genetic divergence (Dxy) for two categories of genes.** Genes with trans-specific polymorphism (yellow) showed higher genetic divergence between species than genes without trans-specific polymorphism (Student’s *t*-test, *p*-value<0.001).

**S4 Fig. Scatterplot of Dxy as function of the coverage depth per transcript for two categories of genes.** Genes with trans-specific polymorphism (yellow) are not associated to higher transcript coverage in greater Dxy values than genes without trans-specific polymorphism.

**S5 Fig. Transcripts z-score distribution from genotypic data.** Based on allele frequencies, each locus score obtained from cRDA was transformed in z-score. Full and dashed red lines correspond to significance threshold of *p*-value of 0.01 and 0.001, respectively.

**S6 Fig. Shift of allele frequencies in outliers SNP identified by multivariate analysis.** A) 30 randomly chosen SNPs from the 348 outliers identified by cRDA showing shift of the minor allele frequencies (MAF) between limnetic and benthic species across continents. 15 randomly chosen SNPs focusing on allele frequencies changes within region highlighting the parallelism of allele frequency shift among regions, in B) Maine (*C. clupeaformis*) C) Norway (*C. lavaretus*) and D) Switzerland (*C. lavaretus*). Each of the 15 SNPs has a different color for comparisons among regions

**S7 Fig. Principal component analysis on gene expression level between individuals.** The PCA indicates the transcriptomic relationships between individuals of Cliff (black), Indian (red), Langfjordvatn (green), Skrukkebukta (lightblue), Zurich (pink) and Lucerne (blue) lakes for limnetic (circles) and benthic (triangles) species are projected along PC1 and PC2. PC1 separated lakes from Norway and Switzerland, but also Cliff and Indian Lakes. PC2 allowed discriminating populations of *C. clupeaformis* form *C. lavaretus*. Indian, Skrukkebukta and Zurich (less differentiated lakes per regions) showed more inter-indivual gene expression level variation. B) MDS projection of the raw individuals counts. Each point corresponds to an individual and each color to a population. Dwarf = Limnetic species; Normal = Benthic species. A_ = America; E_ = Europe. C, I, LF, LU, SK, ZU correspond to Cliff, Indian, Langfjordvatn, Lucerne, Skrukkebukta and Zurich, respectively. No individual clustering (*i.e.*, bias) was associated to sampling location, storage methods, RNA extraction or sequencing.

**S8 Fig. Expression fold changes for differentially expressed genes for intra-lake Limnetic/Benthic species comparisons.** Each density plot is associated to a lake with the corresponding color. Positive and negative fold changes (log2) are associated to an over-expression and an under-expression, respectively, for genes in limnetic species relative to the benthic species.

**S9 Fig. Expression fold changes for differentially expressed genes for intra-region Limnetic/Benthic species comparisons.** Each density plot is associated to a region and composed by overlap between DEGs within region with the corresponding color. Positive and negative fold changes (log2) are associated to an over-expression and an under-expression, respectively, for genes in limnetic species relative to the benthic species.

**S10 Fig. Expression fold changes for differentially expressed genes for across continents Limnetic/Benthic species comparisons.** Each density plot is associated to a continental comparison and composed by overlap between DEGs among regions with the corresponding color. Positive and negative fold changes (log2) are associated to an over-expression and an under-expression, respectively, of genes in limnetic species relative to the benthic species.

**S11 Fig. Distribution of transcripts coverage.** Vertical red lines correspond to the mean level of coverage in DEGs.

**S12 Fig. Transcripts z-score distribution from abundance transcripts data.** Each transcript score obtained from cRDA was transformed in z-score. Full and dashed red lines correspond to significance threshold of *p*-value of 0.01 and 0.001, respectively.

**S13 Fig. Merged significant subnetworks for Limnetic/Benthic species comparisons.** Detailed subset of differentially expressed genes between species showing convergence across the studied system, separately displayed for each metabolic pathway. Gene annotation is based on the Ensembl nomenclature. For each node represented by a gene, the relative size is proportional to the contribution score of the associated gene to the significance of the metabolic pathway. The score for each gene corresponds to a probability of convergent adaptation between individuals of the same species. The color of each node is associated to the fold change expression value. Cold and warm colors correspond to negative and positive fold change values, respectively defined as under- and over-expression in the limnetic species relatively to the benthic species.

**S14 Fig. Significant subnetworks for Limnetic/Benthic species comparisons.** The highest scoring genes within each metabolic pathway are shown in red, composing a subset of significant differentially expressed genes (DEGs), between species. Pyruvate kinase (PKM) gene expression (in yellow) is associated with a *cis*-eQTL in its 3’UTR. Gene identification is based on the Ensembl nomenclature. For each node, the relative size is proportional to the contribution score of the associated gene to the significance of the metabolic pathway. The score for each gene corresponds to a probability of convergent adaptation.

**S15 Fig. Distribution of inferred eQTL p-values.** The p-value of all eQTLs inferred with Matrix eQTL does not show deviation that could be caused by unaccounted covariates. Plot obtained directly from Matrix eQTL.

**S16 Fig. Association between a *cis*-eQTL genotypes and level of expression of the PK gene between species.** Transcripts abundance per individual (circles) as a function of genotypes for the *cis*-eQTL locus. The grey line corresponds to the linear model fitted to the data and showing significant differential expression between genotypes (*P*=0.001) represented species for each continent.

**S1 Table. Number of raw and filtered reads obtained per individual**

**S2 Table. S2 Table. Number of contigs from *de novo* raw assemblies to final used reference, through filtering steps.** For each lineage-specific transcriptome assembly (*i.e., C. lavaretus* and *C. clupeaformis*) and the common reference, number of contigs are reported for the raw assemblies, as well as for the intermediate filtering steps (ORF, longest isoform per transcript and TPM normalization) and uniquely annotated transcripts.

**S3 Table. Gene ontology analysis results for significant enrichment.** Significant GO term (*P*<0.05) from different analyses. From expression data, with identification of DEGs based on DEseq2 (FDR<0.05) (FDR<0.05) and cRDA by applying different thresholds (*P*<P<0.01 and *P*<P<0.001, respectively), and from genotype data from cRDA with a significance threshold of P<0.01.

**S4 Table. Significant subnetworks from the KEGG database.** The significant subnets (*P*<0.05), the number of genes composing the subnet and their respective pathways are indicated. The subnet composition is detailed by the Entrez ID of genes. Aiming to control for any bias in the significance threshold caused by highly expressed genes, we normalized the distribution of the counts.

